# Recovery of Information from Latent Memory Stores Decreases Over Time

**DOI:** 10.1101/537076

**Authors:** Asal Nouri, Edward F. Ester

## Abstract

Working memory (WM) performance can be enhanced by an informative cue presented during storage. This effect, termed a retrocue benefit, can be used to explore how observers prioritize information stored in WM to guide behavior. Recent studies have demonstrated that neural representations of task-relevant memoranda are strengthened following the appearance of a retrocue, suggesting that participants can consult additional information stores to supplement active memory traces. Here, we sought to better understand the nature of these memory store(s) by asking whether they are subject to the same temporal degradation seen in active memory representations during storage. We tested this possibility by reconstructing and quantifying representations of remembered positions from alpha-band EEG activity while varying the interval separating the encoding display and a valid cue in a retrospectively cued spatial WM task. We observed a significant increase in the quality of location-specific representations following a retrocue, but the magnitude of this benefit was linearly and inversely related to the timing of the retrocue such that later cues yielded smaller increases. This result suggests that participants’ ability to supplement active memory representations with information from additional memory stores is not static: the information maintained in these stores may be subject to temporal degradation, or these stores may become more difficult to access with time.

---

Working memory (WM) enables the representation of information no longer present in the sensorium. This system is a core component of general cognitive function, as evidenced by robust correlations between measures of WM ability and aptitudes such as IQ, scholastic achievement, and reading ability (e.g., Daneman & Carpenter, 1980; Engle et al., 1999). Despite this importance, WM is famously burdened with a limited capacity (Luck & Vogel, 2013; Ma et al., 2014). Thus, mechanisms of selective attention are needed to gate access to WM and to prioritize existing WM representations for behavioral output. Attentional prioritization in WM can be studied using retrospective cues (see Souza & Oberauer, 2016; Myers et al. 2017; for recent comprehensive reviews). In a typical retrocue experiment, participants are asked to store an array of objects (e.g., colored squares) for subsequent report. At some point during the ensuing delay period, a cue indicates which of the original memory items is most likely to be tested. Relative to a neutral-cue or no-cue baseline, valid cues yield superior memory performance. This effect is commonly referred to as a retrocue benefit.

Several mechanisms may be responsible for retrocue benefits, including (but not limited to): (a) an attentional enhancement of the cued memory representation (Rerko, Souza, & Oberauer, 2014; Souza, Rerko, & Oberauer, 2015), (b) insulation of the cued memory representation from subsequent decay or interference (Pertzov, Bays, Joseph, & Husain, 2013), (c) removal of non-cued representations from memory (Souza & Oberauer, 2016), or (d) some combination of the above. A recent study (Ester, Nouri, Rodriguez, 2018) sought to evaluate these alternatives by examining changes in neural representations of memoranda following the presentation of a retrocue. Participants were required to remember the spatial positions of two colored discs over a blank delay and report the location of a probed disc via a mouse click. During valid trials, a color retrocue presented during the blank delay informed participants which disc’s location would be probed at the end of the trial. Valid cues were presented either immediately after the offset of the encoding display (valid-early trials) or midway through the subsequent blank delay (valid-late trials). To examine the effects of retro-cues on spatial WM representations, Ester et al. (2018) used an inverted encoding model to reconstruct time-resolved representations of the remembered disc positions from spatiotemporal patterns alpha-band activity (8-12 Hz) recorded over occipitoparietal electrode sites. During neutral trials, the authors observed a monotonic decrease in the quality of reconstructed representations over the course of the delay period. During valid trials, decreases in the quality of the reconstructed representation of the cued disc were either eliminated (valid-early trials) or partially reversed (valid-late trials), while decreases in the quality of the reconstructed representation of the non-cued disc were hastened.

Like others (e.g., Rose et al., 2016; Wolff et al., 2017; Christophel et al., 2018), Ester et al. interpreted the partial recovery in location-specific representations observed during valid-late trials as evidence that participants were able to supplement active memory representations indexed by alpha-band activity with other information sources not indexed by this activity, including but not limited to “activity silent” WM or long-term memory. In this study, we asked whether participants’ ability to exploit these additional memory stores is time-dependent. Specifically, we wondered whether information stored in “offline” memory systems degrades or becomes more difficult to access over time, much in the same way that active memory representations indexed by alpha-band activity degrade during neutral trials (Ester et al., 2018). If so, then one would expect that the extent of retrocue-driven recovery in reconstructed representations would decrease as the temporal interval separating the encoding display and the retrocue increases. We tested this possibility by replicating the neutral and valid-late conditions from Ester et al. (2018) while varying the timing of the retrocue between 1.0, 1.5, and 2.0 sec after termination of the encoding display. As expected, we found that degree of recovery in reconstructed representations of position was linearly and inversely related to the interval separating the retrocue and the encoding display, suggesting that position information stored in “offline” memory systems may degrade or become more difficult to access over time.

## Methods

### Participants

Thirty-six (36) volunteers were recruited from the Florida Atlantic University undergraduate community and tested in a single 2.5-hour session. All experimental procedures were approved by the local institutional review board. Participants gave both written and oral informed consent, self-reported normal or corrected-to-normal visual acuity, and were compensated with course credit or monetary remuneration ($15/h in Amazon gift cards).

### Testing Environment

Participants were tested in a dimly lit, sound-attenuated recording chamber. Stimulus displays were generated in MATLAB and rendered on a 17-inch dell CRT monitor cycling at 85 Hz (1024 x 768 pixel resolution) via Psychophysics Toolbox software extensions (Kleiner et al., 2007). Participants were seated approximately 60 cm from the display (head position was not constrained) and made responses using a standard computer keyboard and mouse.

### Behavioral Task

A trial schematic is shown in Figure 1. Participants were instructed to fixate a small dot (subtending 0.2° from a viewing distance of 60 cm) throughout the experiment. Each trial began with a sample display containing two colored discs (one red and one blue). Each disc was presented in one of 8 equally spaced positions (22.5° to 337.5° in 45° increments) along the perimeter of an imaginary circle (radius 6° visual angle) centered at the fixation point (Figure 1B). A small amount of jitter (±10° polar angle) was added to the location of each disc on each trial to discourage verbal coding strategies (e.g., “the blue disc was at 2 o’clock”).

**Figure 1.**
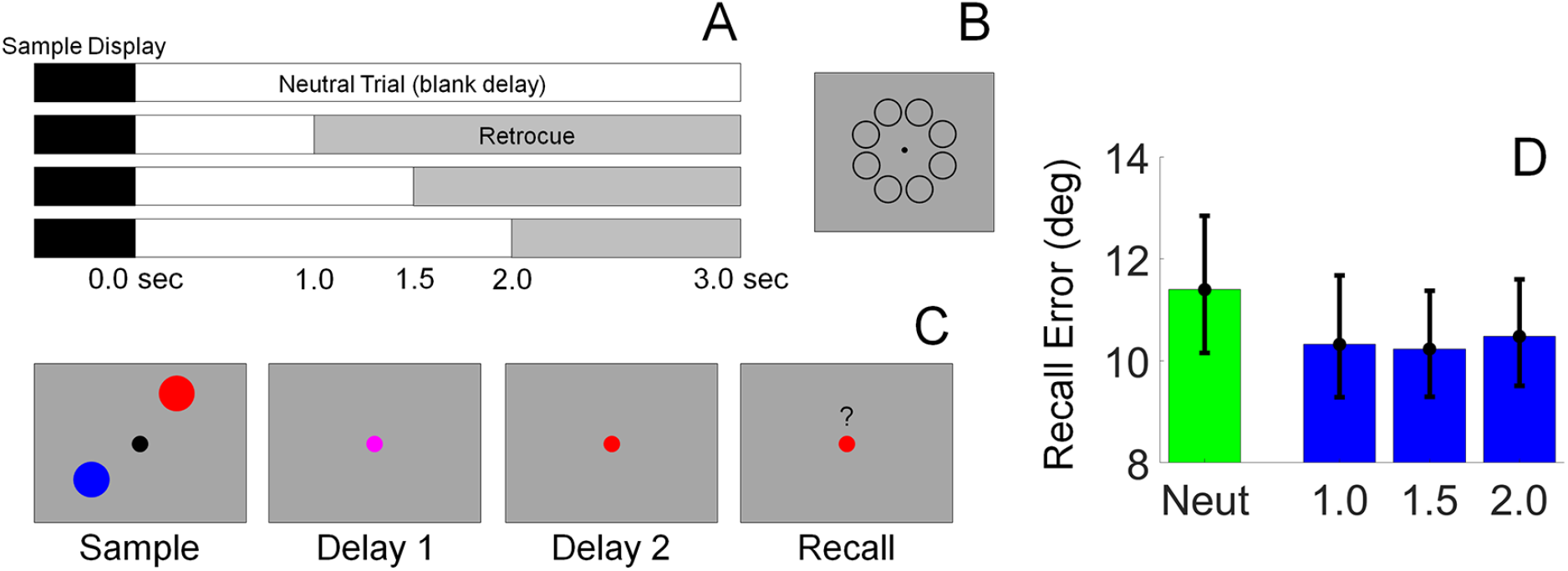
Task Design and Behavioral Data. (A) Schematic showing the different cue conditions. On 50% of trials, the sample display was followed by a 3.0 sec blank delay (neutral trials). On the remaining 50% of trials, a 100% valid retrocue was rendered 1.0, 1.5, or 2.0 sec after termination of the sample display. In all cases, the total delay period length was fixed at 3.0 seconds. (B) Schematic showing possible disc positions. (C) Schematic showing the sequence of events during each trial (Note: stimuli are not drawn to scale; see Methods for information regarding display and stimulus geometry). (D) Behavioral performance, quantified as average absolute recall error (i.e., the average absolute difference between the polar location reported by the participant and the polar location of the probed disc) Error bars depict the 95% confidence interval of the mean.

The sample display was extinguished after 0.5 sec, at which point the fixation point changed colors from black to purple. The fixation point either remained purple for the ensuing 3.0 sec delay period (neutral trials) or changed colors from purple to either blue or red 1.0, 1.5, or 2.0 sec after termination of the sample display (valid trials). When presented, valid cues indicated which disc participant would be probed to report at the end of the trial with 100% reliability. Each trial concluded with a test display containing a blue or red fixation point, a mouse cursor, and a question mark symbol (“?”) above the fixation point. Participants were required to click on the location of the disc indicated by the color of the fixation point within a 3.0 sec response window. Memory performance was quantified as the absolute angular distance between the polar location of the probed disc and the polar location reported by the participant. Performance feedback was given at the end of each block. Participants completed 6 (N = 1), 10 (N = 1), 11 (N = 1) or 12 (N = 33) blocks of 56 trials.

### EEG Recording and Preprocessing

Continuous EEG was recorded from 63 Ag/Ag-Cl^−^ scalp electrodes via a Brain Products actiCHamp amplifier. An additional electrode was placed over the right mastoid. Data were recorded with a right mastoid reference and later re-referenced to the algebraic mean of the left and right mastoids (10-20 site TP9 served as the left mastoid reference). The horizontal and vertical electrooculogram (EOG) was recorded from electrodes placed on the left and right canthi and above and below the right eye, respectively. All electrode impedances were kept below 15 kΩ, and recordings were digitized at 1000 Hz. Recorded data were bandpass filtered from 1 to 50 Hz (3^rd^ order zero-phase forward and reverse Butterworth filters), epoched from a period spanning −1000 to +4500 ms relative to the start of each trial, and baseline corrected from −250 to 0 ms. Muscle and electrooculogram artifacts were removed from the data using independent components analysis (ICA) as implemented in EEGLAB (Delorme & Makeig, 2004).

### Inverted Encoding Model

Following earlier work (e.g., Foster et al. 2016; Ester et al. 2018) we used an inverted encoding model to reconstruct location-specific representations of the red and blue discs during the sample display and subsequent delay period. Reconstructions of stimulus locations were computed from the spatial topography of induced alpha-band (8-12 Hz) power measured across 17 occipitoparietal electrode sites: O1, O2, Oz, PO7, PO3, POz, PO4, PO8, P7, P5, P3, P1, Pz, P2, P4, P6, and P8. To isolate alpha-band activity, the raw EEG time series at each electrode was bandpass filtered from 8-12 Hz (zero-phase forward and reverse filters as implemented by EEGLAB’s “eegfilt” function; 24 dB/octave attenuation), yielding a real-valued signal *f*(*t*). The analytic representation of *f*(*t*) was obtained by applying a Hilbert transformation:

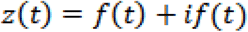

where *i* = √−1 and *if*(t) = *A*(*t*)*e*^*iφ*(*t*)^. Induced alpha power was computed by extracting and squaring the instantaneous amplitude *A*(*t*) of the analytic signal *z*(*t*).

We modeled alpha power at each scalp electrode as a weighted sum of 8 location-selective channels, each with an idealized tuning curve (a half-wave rectified cosine raised to the 8^th^ power). The maximum response of each channel was normalized to 1, thus units of response are arbitrary. The predicted responses of each channel during each trial were arranged in a *k* channel by *n* trials design matrix *C*. Separate design matrices were constructed to track the locations of the blue and red discs across trials (i.e., we reconstructed the locations of the blue and red discs separately, then later sorted these reconstructions according to cue condition).

The relationship between the data and the predicted channel responses *C* is given by a general linear model of the form:

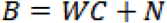

where B is a *m* electrode by *n* trials training data matrix, W is an *m* electrode by *k* channel weight matrix, and *N* is a matrix of residuals (i.e., noise).

To estimate *W*, we constructed a “training” data set containing an equal number of trials from each stimulus location (i.e., 45-360° in 45° steps) condition. We first identified the location φ with the fewest *r* repetitions in the full data set after EOG artifact removal. Next, we constructed a training data set *B_trn_* (*m* electrodes by *n* trials) and weight matrix *C_trn_* (*n* trials by *k* channels) by randomly selecting (without replacement) 1:*r* trials for each of the eight possible stimulus locations (ignoring cue condition; i.e., the training data set contained a mixture of neutral and valid trials). The training data set was used to compute a weight for each channel *C_i_* via least-squares estimation:

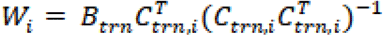

where *C_trn,i_* is an *n* trial row vector containing the predicted responses of spatial channel *i* during each training trial.

The weights *W* were used to estimate a set of spatial filters *V* that capture the underlying channel responses while accounting for correlated variability between electrode sites (i.e., the noise covariance; Kok et al. 2017):

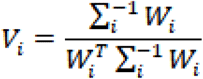

where *Σ_i_* is the regularized noise covariance matrix for channel *i* and estimated as:

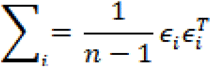

where *n* is the number of training trials and *ε_i_* is a matrix of residuals:

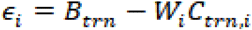

Estimates of ε_*i*_ were obtained by regularization-based shrinkage using an analytically determined shrinkage parameter (see Blankertz et al. 2011; Kok et al. 2017). An optimal spatial filter *v_i_* was estimated for each channel *C_i_*, yielding an *m* electrode by *k* filter matrix *V*.

Next, we constructed a “test” data set *B_tst_* (*m* electrodes by *n* trials) containing data from all trials not included in the training data set and estimated trial-by-trial channel responses *C_tst_* (*k* channels x *n* trials) from the filter matrix *V* and the test data set:

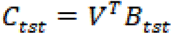

Trial-by-trial channel responses were interpolated to 360°, circularly shifted to a common center (0°, by convention), and sorted by cue condition (e.g., neutral vs. valid). To quantify the effects of retrospective cues reconstructed representations, we obtained an estimate of representation strength by converting the (averaged) channel response estimates for each cue condition to polar form and projected them onto a vector with angle 0°:

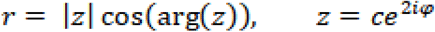

where *c* is a vector of estimated channel responses and *φ* is the vector of angles at which the channels peak.

To ensure internal reliability this entire analysis was repeated 50 times, and unique (randomly chosen) subsets of trials were used to define the training and test data sets during each permutation. The results were then averaged across permutations.

### Eye Movement Control Analyses

We used independent component analysis to isolate and remove electrooculogram and muscle artifacts from the data. Nevertheless, small but reliable biases in eye position towards the location(s) of the remembered disc(s) could contribute to reconstructions of stimulus location. We tested this possibility by regressing trial-by-trial horizontal EOG recordings (in μV) onto the horizontal position of the cued disc (see Foster et al., 2016). We restricted our analysis to valid trials on the assumption that eye movements would be most prevalent when participants were explicitly told which disc would be probed. In this framework, positive regression coefficients reflect greater changes in eye position as a function of stimulus location. Separate regressions were run for each participant and the resulting coefficients were averaged across participants.

### Statistical Comparisons

All statistical comparisons were based on non-parametric permutation (bootstrap) tests. For example, to compare the magnitude of the retrocue benefit during the 1.0 and 1.5 cue delay conditions, we randomly selected (with replacement) and averaged data from 36 of 36 participants in each condition, then computed the difference between the two means. This procedure was repeated 10,000 times, yielding a 10,000-element distribution of average differences. Finally, we computed the number of permutations where the cue benefit was opposite the expected direction, yielding an empirical p-value. Where appropriate, p-values were false-discovery-rate (FDR) corrected for multiple comparisons using the Benjamini & Hochstein (1995) procedure. Effect sizes for key contrasts were computed using a nonparametric method 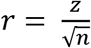, where *z* is the outcome of a Wilcoxon signed rank test (Rosenthal, 1994).

### Within-participant Variability

We report estimates of within-participant variability (e.g., 95% within-participant confidence intervals) throughout the paper. These estimates discard subject variance (e.g., overall differences in response strength) and instead reflect variance related to the subject by condition(s) interaction term(s) (e.g., variability in response strength across experimental conditions; Loftus & Masson, 1994; Cousineau 2005). We used the approach described by Cousineau (2005): raw data (e.g., location information or channel response estimates) were de-meaned on a participant by participant basis, and the grand mean across participants was added to each participant’s zero-centered data. The grand mean centered data were then used to compute bootstrapped within-participant confidence intervals (10,000 permutations).

## Results

We recorded EEG while participants performed a retrospectively cued spatial WM task (Figure 1A-C). Participants were required to remember the polar locations of two colored discs over a blank delay period. After termination of the sample display the fixation point changed colors from black to purple, instructing participants to remember the positions of both discs. During neutral trials the fixation point remained purple throughout the delay period, then changed colors from purple to either blue or red during the recall epoch. This color change served as a prompt for participants to report the position of the disc whose color matched the fixation point via a mouse click. During valid trials, the fixation point changed from purple to either blue or red 1.0, 1.5, or 2.0 seconds following termination of the sample display. This color change informed participants which disc they would be probed to report at the end of the trial. Participants’ behavioral performance was quantified as mean absolute recall error, or the average mean absolute difference between the polar angle reported by the participant and the polar location of the probed disc. Report errors were reliably lower during all three cue conditions relative to neutral trials (false-discovery-rate-corrected bootstrap tests, *p* < 0.006), but errors did not differ across valid cue conditions (1.0, 1.5, and 2.0 s; FDR-corrected p-values > 0.27; Figure 1D). These findings are consistent with earlier reports suggesting that valid retrospective cues can improve memory performance (e.g., Griffin & Nobre, 2003; Sprague et al. 2016; Ester et al. 2018).

Following earlier studies (Foster et al., 2016; Ester et al., 2018), we used a spatial inverted encoding model to recover a representation of each disc’s polar position from spatiotemporal patterns of occipitoparietal alpha-band power. Specifically, we modeled induced alpha power recorded at each electrode site as a weighted sum of eight location-selective channels, each with an idealized response function. The resulting weights were used to calculate a predicted response for each channel given patterns of induced alpha power recorded during each trial of a statistically independent data set. Trial-by-trial reconstructions were circularly shifted to a common center (0° by convention), yielding a single time-resolved 360° channel response function for each participant (Figure 2). We quantified these reconstructions by converting them to polar form and projecting them onto a unit vector with angle 0°. The resultant vector length was interpreted as a measure of total location information.

**Figure 2.**
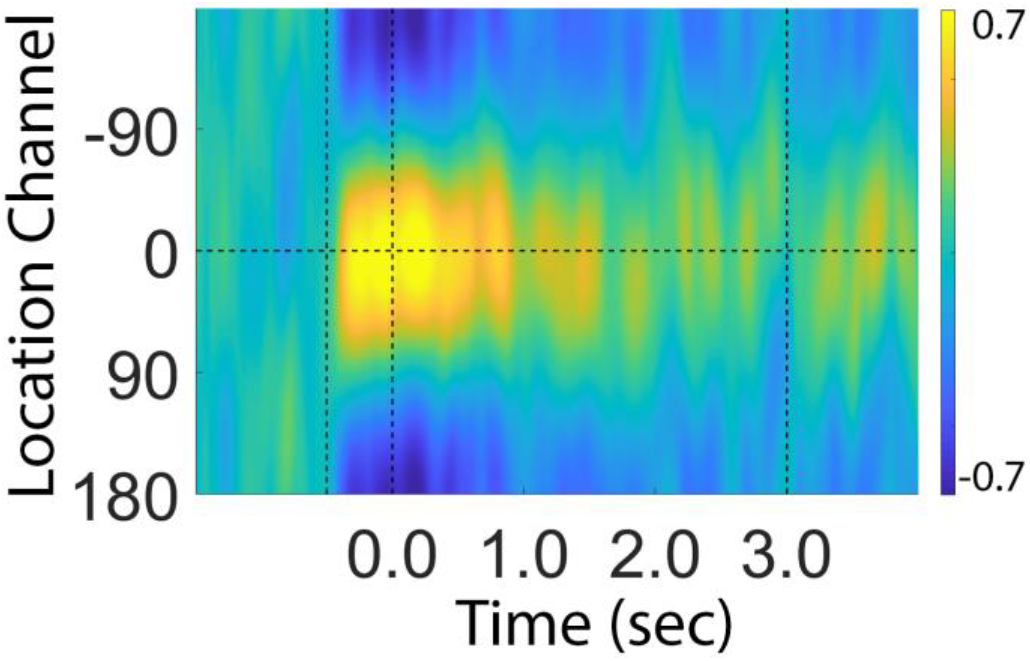
Average Reconstructed Channel Responses During Neutral Trials. Dashed vertical bars at 0.0 and 3.0 s denote the start and end of the memory delay. Participant-level reconstructions were converted to polar form and projected onto a vector with angle 0°. The resulting vector length was interpreted as a measure of total location information (e.g., Figure 3).

As shown in Figure 3, during neutral trials estimates of representation strength declined monotonically throughout the delay period (−21.04 units/sec; p < 0.0001). However, visual inspection suggests that this decline was partially reversed during valid trials. We tested this possibility by comparing average location information for the cued disc during valid trials across epochs spanning 0-500 ms immediately after presentation of the cue and near the end of the trial (2500-3500 ms after termination of the sample display). The 0-500 ms after-cue epoch was selected based on studies suggesting that it takes participants approximately 400 to 600 ms to process and utilize a retrospective cue (e.g., Pertzov et al., 2013), while the 2500-3500 ms end-of-trial epoch was selected to allow sufficient time for recovery during the 2.0 s cue condition^1^. If presentation of a valid retrocue enables a partial recovery of location information (e.g., from latent memory sources), then estimates of location information should be reliably higher during the end-of-trial relative to the after-cue epoch. This was true during the 1.0 s cue condition (Figure 4; M = −3.49 vs. 77.34 units for the after-cue and end-of-trial epochs; FDR-corrected p-value = 0.008, r = 0.44), but not during the 1.5 s or 2.0 s cue conditions (p > 0.15). These findings are broadly consistent with those reported by Ester et al. (2018): memory representations reconstructed from alpha-band EEG activity degrade with time, but a valid retrocue can partially reverse earlier degradation.

**Figure 3.**
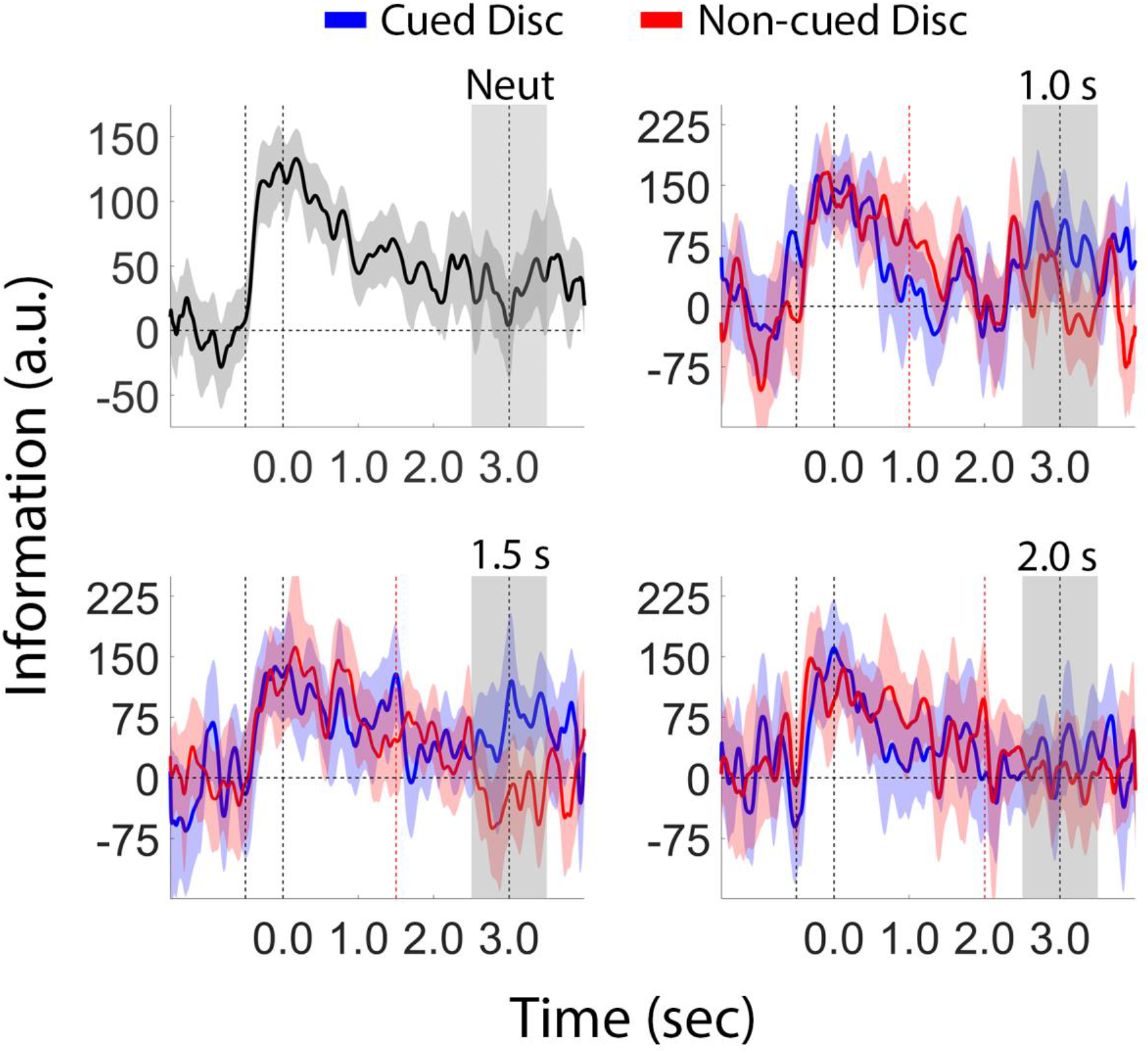
Estimates of Location Information Recovered from EEG Activity. Black vertical dashed bars at 0.0 and 3.0 sec indicate the start and end of the blank delay period. Red vertical dashed bars indicate the onset of the retrocue during valid trials. The shaded region from 2.5 to 3.5 seconds was used in later statistical comparisons (see Figures 3 and 4). The shaded area around each plot depicts the 95% within-participant confidence interval of the mean.

**Figure 4.**
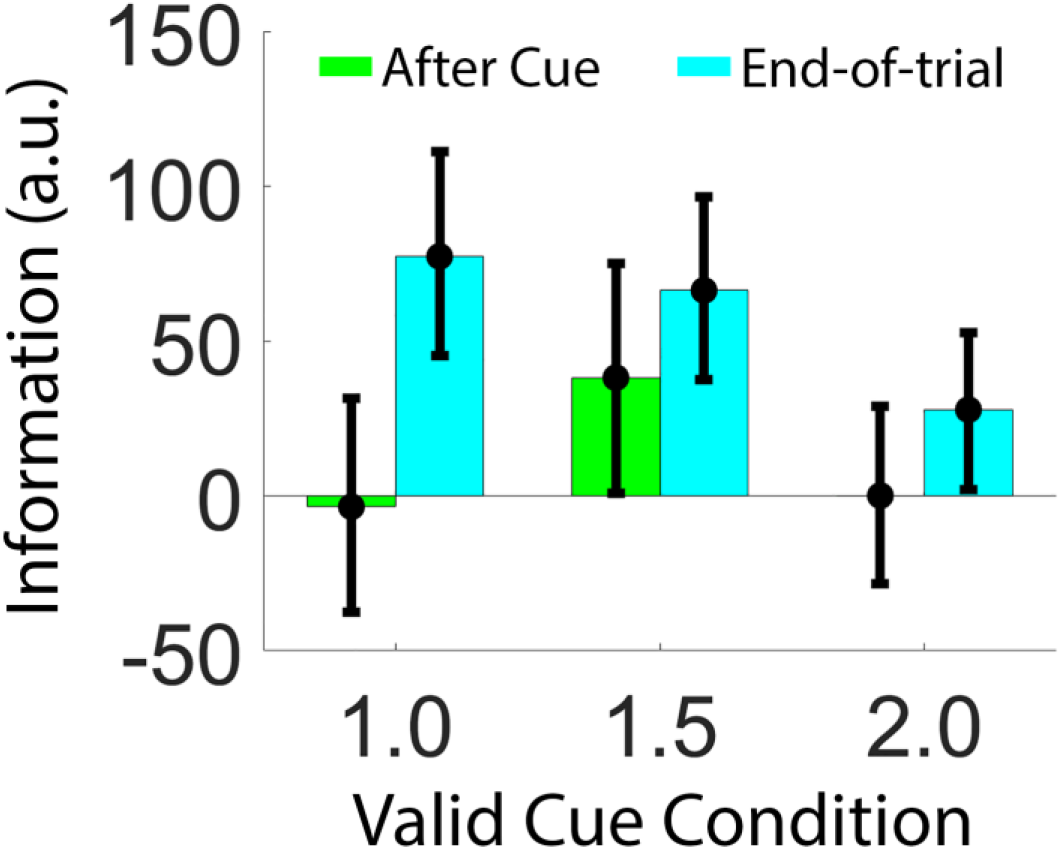
Recovery of Information Following a Retrocue. Time-based changes in location information for the cued disc during periods spanning 0-500 ms following cue onset (“after cue”) and 2500 to 3500 ms after termination of the sample display (“end-of-trial”). Error bars depict the 95% within-participant confidence interval of the mean.

### The recovery of information following a retrospective cue declines with time

Recovery of information following a retrocue suggests that participants can supplement active memory representations with information stored in memory systems not indexed by alpha-band activity, such as activity-silent short-term memory or long-term memory. The goal of this study was to examine whether information held in these alternative stores degrades or becomes more difficult to access over time. If so, then the amount of information recovery seen in representations of memoranda reconstructed from alpha band activity (Figures 3 and 4) should decrease as the temporal interval separating the encoding display from the retrocue increases. We tested this possibility by calculating the total retrocue benefit as a function of cue timing. We defined the total retrocue benefit as the difference in information between the cued disc during valid trials and the average of the two remembered discs during neutral trials over an interval spanning 2.5 to 3.5 sec following termination of the sample display. These estimates are plotted in Figure 5. Significant retrocue benefits were observed during the 1.0 and 1.5 s valid cue conditions, but not during the 2.0 s condition (M = 44.84, 33.85, and −4.64 units; p = 0.01, 0.04, and 0.62, respectively). The pattern across cue conditions was well-approximated by a linear trend with a significant negative slope (−49.58 units/sec; p = 0.009; Figure 5B), consistent with the hypothesis that participants’ ability to exploit information retained in other memory stores is time-limited.

**Figure 5.**
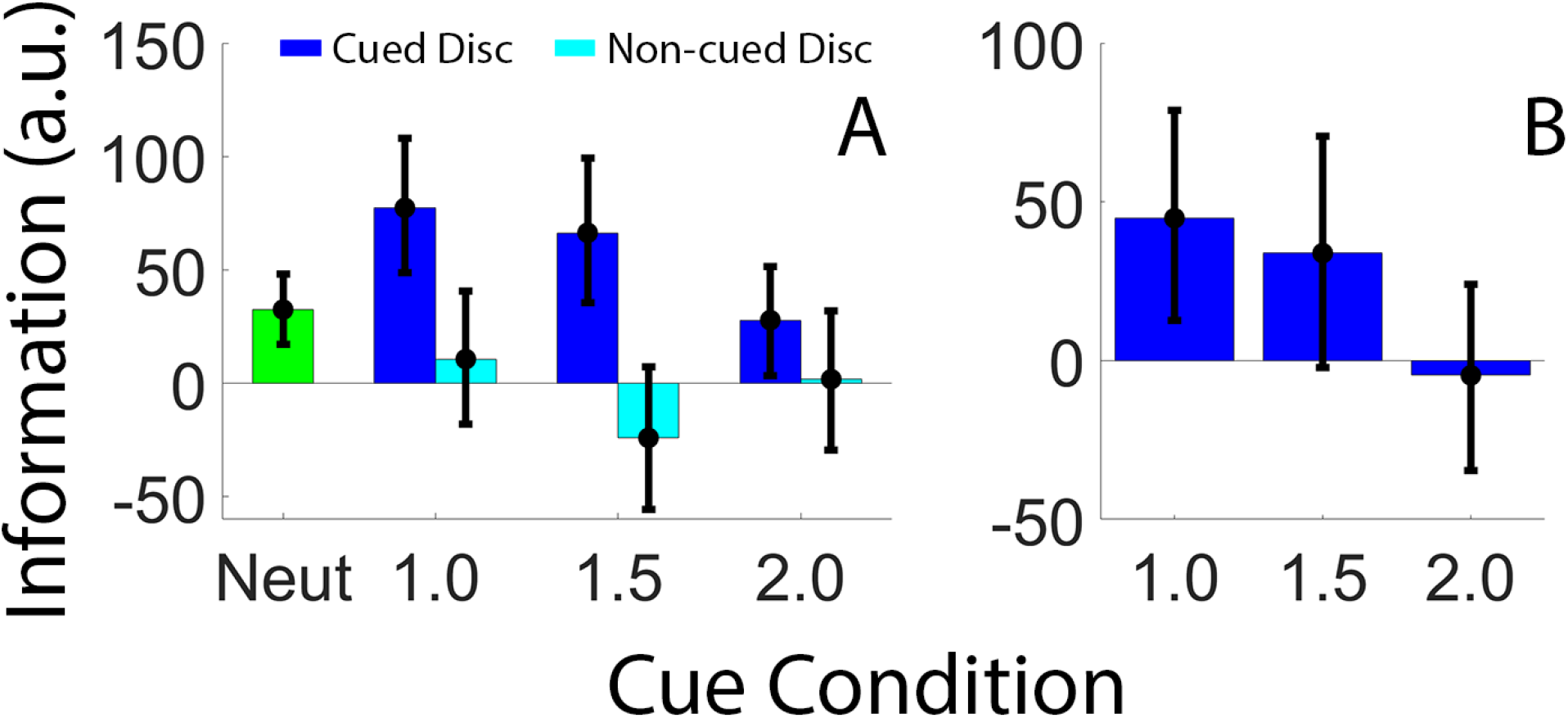
Retrocue Benefits Decrease with Time. (A) Location information estimates for the cued and non-cued disc averaged across an epoch spanning 2500-3500 ms after termination of the sample display. Information estimates were averaged across both remembered discs during neutral trials (green bar). (B) Total retrocue benefit across valid cue conditions, defined as the difference between location information for the cued disc minus average location information for both discs during neutral trials. Error bars depict the 95% within-participant confidence interval of the mean.

### Ruling out contributions from eye movements

We identified and removed electrooculogram artifacts from the data via independent components analysis. However, small and consistent eye movement patterns opaque to ICA could nevertheless contribute to the location reconstructions reported here. We examined this possibility by regressing time-resolved estimates of horizontal EOG activity onto remembered stimulus locations. As shown in Figure 6, the regression coefficients linking eye position with remembered locations were indistinguishable from 0 for the duration of each trial, suggesting that eye movements were not a major determinant of location reconstructions.

**Figure 6.**
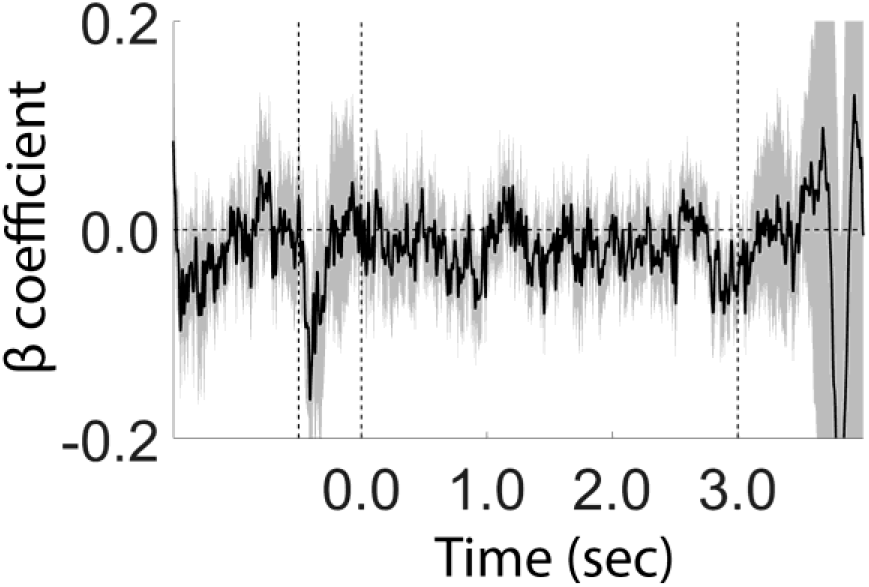
Effect of Eye Position on Stimulus Reconstructions. Black vertical dashed lines at 0.0 and 3.0 depict the start and end of the delay period. Shaded regions depict the 95% within-participant confidence interval of the mean.

### Location Information is Absent in Other Frequency Bands and Electrode Sites

Extant data suggests that posterior alpha band power indexes the focus of covert spatial attention and the contents of spatial working memory while activity in other frequency bands does not (e.g., Foster et al., 2016; Foster et al., 2017). For example, Foster et al. (2016) attempted to reconstruct a representation of a remembered position by applying an inverted encoding model to induced power in multiple frequency bands (4-50 Hz) and found that robust stimulus reconstructions could only be obtained using activity between approximately 7-14 Hz. Our data accord with this conclusion. We applied an inverted encoding model to theta-band (4-7 Hz) and beta-band (13-30 Hz) recorded over the same occipitoparietal electrodes used in our primary analysis (we did not model gamma band activity as there is active debate regarding whether high frequency [> 30 Hz] activity measured with EEG indexes cortical dynamics or electromyogenic artifacts; e.g., Muthukumaraswamy, 2013; Nunez & Srinivasan, 2010; Yuval-Greenberg et al., 2008). As shown in Figure 7, we were unable to recover reliable location information from activity in either theta or beta band activity in any experimental condition. Thus, consistent with earlier work, position-specific signals that enable decoding and reconstruction of attended or remembered positions appear to be uniquely indexed by alpha band activity.

**Figure 7.**
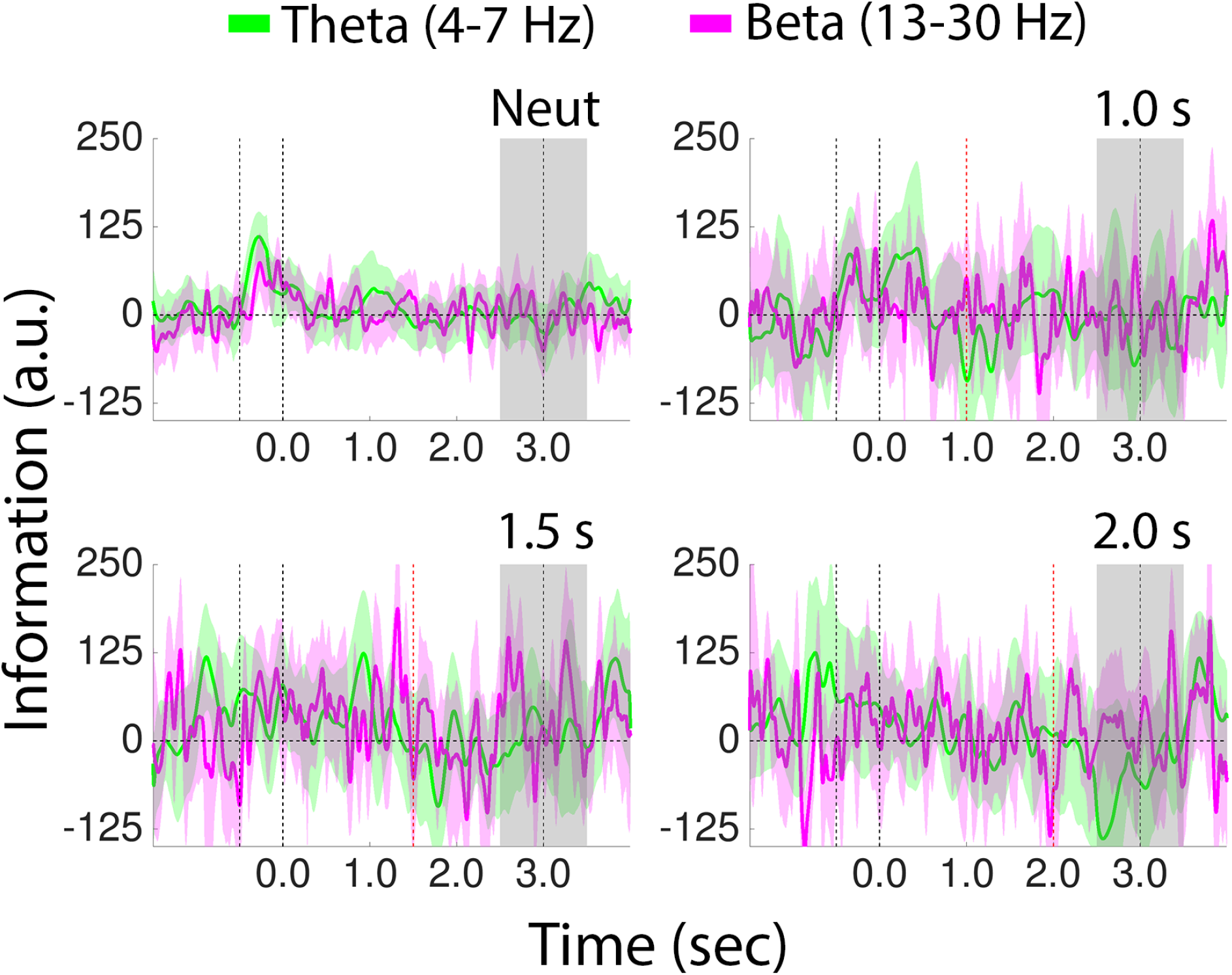
Location Information Cannot Be Recovered from Posterior Theta and Beta Band Activity. Data shown for the 1.0, 1.5, and 2.0 s valid conditions reflect location information for the cued disc only, while data shown for the neutral condition reflect average location information for the two remembered discs. Shaded regions depict the 95% within-participant confidence interval of the mean.

Executive functions such as attention and working memory have long been associated with activity in prefrontal cortex. However, it is unclear whether alpha band activity recorded over frontal electrode sites also supports recovery of location information. To test this possibility, we applied the same inverted encoding model used in previous sections to alpha band activity recorded from 12 frontocentral electrodes (10-20 sites Fz, F1, F2, F3, F4, F5, FC1, FC2, FC3, FC4, FC5, and FC6). As shown in Figure 8, we were unable to recover robust estimates of location information over these sites, indicating that position-specific signals enabling the recovery of robust location information are uniquely indexed by posterior (occipitoparietal) alpha band activity.

**Figure 8.**
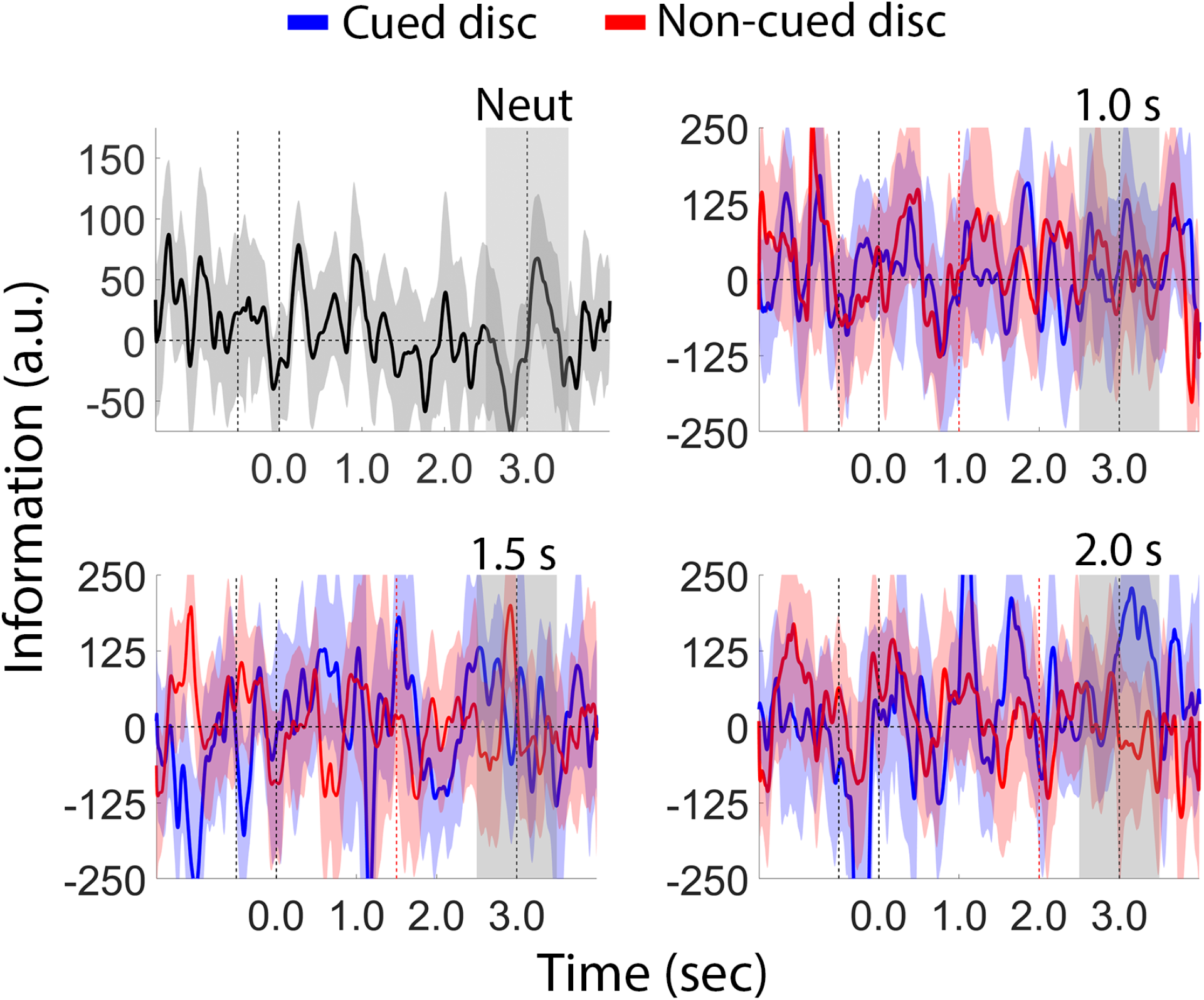
Location Information Cannot Be Recovered from Frontocentral Alpha Band Activity. Conventions are as in Figure 2.

## Discussion

Working memory performance can be improved by an informative retrocue presented during storage. This effect, termed a retrocue benefit, can be used to study mechanisms of attentional priority in WM. Recent neuroimaging (EEG and fMRI) studies have identified a partial recovery in neural representations of cued memory items following a retrospective cue (Sprague et al., 2016; Rose et al., 2016; Wolff et al., 2017; Christophel et al., 2018; Ester et al., 2018), suggesting that participants can query information held in other memory stores to supplement active memory representations. Here, we asked whether these additional stores are subject to the same degradation seen for active representations (e.g., Figure 3). To test this possibility, we used an inverted encoding model to recover and quantify the fidelity of location-specific representations from EEG activity recorded while participants performed a retrospectively cued spatial WM task. In addition, we systematically varied the timing of the retrocue relative to the termination of the encoding display from between 1.0 to 2.0 seconds. Consistent with earlier findings (e.g., Ester et al., 2018) we observed a recovery in location information estimates following the appearance of a retrospective cue (Figures 3–4). However, the magnitude of the retrocue benefit was inversely and linearly related to the interval separating the termination of the encoding display and the onset of the cue (i.e., lower benefits for longer intervals; Figure 5). This result is consistent with the hypothesis that information stored in offline memory systems is subject to temporal degradation (e.g., due to decay or inter-item interference), or that it becomes more difficult to access over time.

A growing body of evidence suggests that human observers can supplement active memory representations with other information sources, including but not limited to “activity silent” WM or long-term memory (e.g., Rose et al., 2016; Sprague et al., 2016; Wolff et al., 2017; Ester et al., 2018; Sutterer et al., 2018). On the one hand, there is ample evidence that long-term memory systems can be used to complement or supplement active memory storage when the task allows (e.g., Carlisle et al., 2011). However, behavioral studies have found little to no evidence for time-based degradation in visual long-term memory. For example, in some instances recall of visual features from long-term memory has been found to match performance during recall from WM (Brady et al., 2008; Brady et al., 2013). Based on these findings, we are skeptical that retrieval from long-term memory can account for the data reported here. On the other hand, models of activity-silent working memory information propose that task-relevant representations are maintained through short-term synaptic plasticity (Zucker & Regehr, 2002). An influential modeling study (Mongillo et al., 2008) suggests that calcium kinetics in prefrontal cortex could act as a short-term memory buffer with a time constant of about 1-2 seconds. In the absence of continuous spiking input, representations encoded in this format would be expected to degrade with time. This prediction is supported by the pattern of findings reported here.

Our analyses focused on posterior (occipitoparietal) alpha band power as extant work suggests that location-specific signals are uniquely indexed by induced activity in the 7-13 Hz range. For example, posterior alpha-band activity covaries with the deployment of spatial attention (e.g., Thut et al., 2006; Rihs et al., 2007), and horizontal and vertical shifts of attention can be decoded from changes in posterior alpha-band power (e.g., Bahramisharif et al., 2010). More recent evidence suggests that posterior alpha-band activity tracks both the locus of covert spatial attention (Foster et al., 2017) and the contents of spatial working memory (Foster et al., 2016). Other frequencies do not appear to share these properties. Like others (Foster et al., 2016) we were unable to recover stimulus position estimates from induce theta- and beta-band activity recorded over occipitoparietal sites (Figure 7). We were also unable to recover stimulus position estimates from alpha-band activity recorded over frontocentral electrode sites (Figure 8). These findings strongly suggest that position-specific signals that enable tracking of covert spatial attention and spatial working memory storage are uniquely indexed by spatiotemporal patterns of alpha band activity recorded over posterior (occipitoparietal) electrode sites.

## Author Contributions

A.N. and E.F.E. conceived and designed the experiment. A.N. collected data; A.N. and E.F.E. analyzed data. A.N. and E.F.E. wrote the paper.

## Acknowledgements

Funding for this project was provided by a State University System of Florida startup award to E.F.E. The authors wish to thank Brian Escobar for assistance with data collection.

## Footnotes

1 The end-of-trial epoch from 2500-3500 ms encompasses portions of the blank memory delay and recall displays. However, we note that physical stimulation during these periods was virtually identical (Figure 1C). Moreover, the recall display contained no additional spatial information that could be used to supplement mnemonic representations of the to-be-recalled disc’s position.

